# The Variability Puzzle in Human Memory

**DOI:** 10.1101/235127

**Authors:** M.J. Kahana, E.V. Aggarwal, T.D. Phan

## Abstract

Memory performance exhibits a high level of variability from moment to moment. Much of this variability may reflect inadequately controlled experimental variables, such as word memorability, past practice and subject fatigue. Alternatively, stochastic variability in performance may largely reflect the efficiency of endogenous neural processes that govern memory function. To help adjudicate between these competing views, we conducted a multisession study in which subjects completed 552 trials of a delayed free-recall task. Applying a statistical model to predict variability in each subject’s recall performance uncovered modest effects of word memorability, proactive interference, and other variables. In contrast to the limited explanatory power of these experimental variables, performance on the prior list strongly predicted current list recall. These findings suggest that endogenous factors underlying successful encoding and retrieval drive variability in performance.

---

Since the first half of the 20th century, students of both human and animal memory have assiduously adhered to the doctrine of functionalism (Carr, 1931; McGeoch, 1942; Woodworth, 1938). This doctrine asserts that we may best arrive at a deeper understanding of memory through the analysis of the functional relations between independent variables, such as similarity or repetition, and dependent variables, such as accuracy and reaction time. With the advent of cognitive neuroscience, functionalists now have a greatly expanded array of dependent variables, including diverse measures of brain function.

Although functionalists have enjoyed considerable success in their efforts to understand memory, the substantial inter-trial variability that is ubiquitous across memory paradigms has limited their progress. Every new student of memory is taught to reduce variability by randomizing or counterbalancing experimental conditions, and designing lists of items matched on word frequency, word length, semantic similarity, and emotional valence, among other measures. Yet, despite our best efforts, performance under similar conditions can vary dramatically from list to list, and from day to day.

The present work seeks to characterize stochastic variability in human memory through a detailed analysis of performance in the classic free recall task. In this task, subjects study lists of common words and then later attempt to freely recall those words in any order they wish. Our basic unit of analysis is the percentage of words remembered following the study of a given list. To examine list-level variability both within and across sessions we recruited 79 subjects for a 23-session experiment in which they contributed data from 552 study-test lists (24 lists per session × 23 sessions). Between the study period and the recall test, subjects performed 24 seconds of a mental arithmetic task. This was done to attenuate recency-sensitive contributions to recall.

## Methods

The data reported here come from Experiment 4 of the Penn Electrophysiology of Encoding and Retrieval Study (PEERS). PEERS aims to assemble a large database on the electrophysiological correlates of memory encoding and retrieval. Although data from Experiments 1-3 have been previously reported (Healey, Crutchley, & Kahana, 2014; Healey & Kahana, 2014, 2016; Lohnas & Kahana, 2013, 2014; Lohnas, Polyn, & Kahana, 2015), this is the first paper reporting data from Experiment 4. Subjects consisted of young adults (ages 18-35) recruited from among the students and staff at the University of Pennsylvania and neighboring institutions.

During each of 23 experimental sessions, subjects (*n* = 79) studied lists of 24 session-unique English words. Immediately following the presentation of each list, subjects performed an arithmetic distractor task for 24 seconds, and then were given 75 seconds to freely recall as many items as they could remember from the just-presented list (delayed free recall).

For each list within a session, words were drawn without replacement from a pool of 576 common English words. To create our word pool for this study, we selected words from among the 1638 word pool used in prior PEERS experiments. The selection process involved removal of words with extreme values along dimensions of word frequency, concreteness, and emotional valence, with the goal of creating a relatively homogenous word pool. Each of these 576 words appeared exactly once in each experimental session (24 lists = 24 items). Within each session, words were randomly assigned to lists following certain constraints on semantic similarity, as described in our earlier PEERS papers^1^.

Each word appeared individually for 1600 ms, followed by an inter-stimulus interval of 800–1200 ms (uniform distribution). Following the presentation of the last word in each list, participants performed a distractor task for 24 seconds. The distractor task consisted of answering math problems of the form *A* + *B* + *C* =?, where *A, B,* and *C* were positive, single-digit integers, though the answer could have been one or two digits. When a math problem was presented on the screen, the participant typed the sum as quickly as possible. Participants were given a monetary bonus based on the speed and accuracy of their responses. After the post-list distractor task, there was a jittered delay of 1200–1400 ms, after which, a tone sounded, a row of asterisks appeared, and the participant was given 75 seconds to freely recall the studied items.

For a random half of the lists in each session (excluding list 1), participants also completed a pre-list distractor task for 24 seconds before presentation of the first word, with a 800-1200 ms (uniform distribution) jittered delay between the last distractor problem and the presentation of the first list word. Subjects were given a short break (approximately 5 minutes) after every 8 lists in a session, which we call a block.

All previously published raw behavioral data from the PEERS studies, as well as the new data reported in the present manuscript, may be freely obtained from the authors’ website, http://memory.psych.upenn.edu.

## Results

We first sought to quantify variability in recall performance across both sessions and lists. In these and subsequent analyses, our basic unit of analysis is the percentage of words remembered following the study of a given list. Figure 1A shows within-subject variability in recall across sessions. For each subject, sorted from lowest to highest on the basis of their overall performance, we marked the average recall for each of their 23 sessions. The vertical dispersion of the dots (each representing the average of 24 lists) illustrates the session-level variability of each subject.

**Figure 1.**
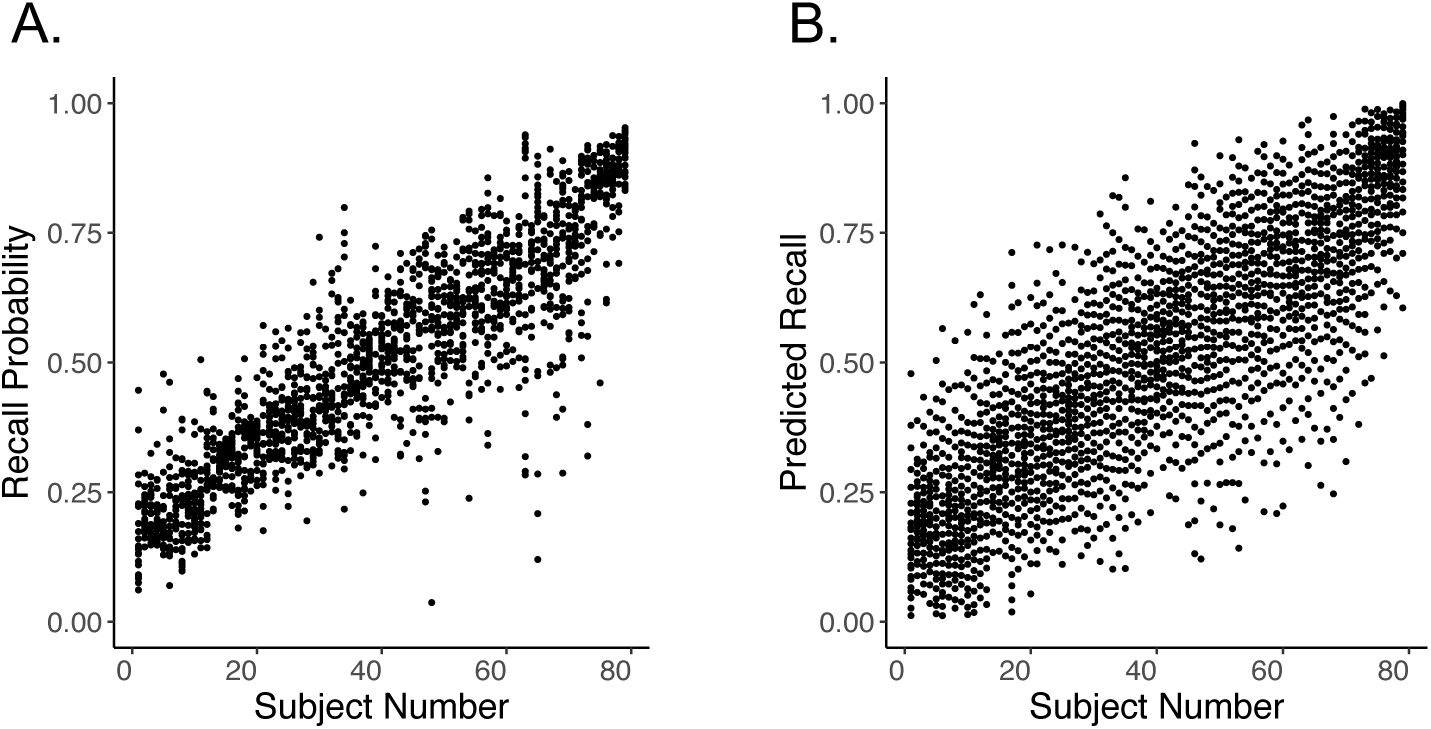
Variability in free recall. **A. Inter-session variability.** Each dot represents the proportion of words recalled for a single subject in a single session. **B. Inter-list variability.** Each dot here represents the proportion of words recalled for a single subject on 23 lists, one list taken from each session, ranked by within-session recall performance.

To illustrate within-session inter-list variability, we again sorted subjects by their overall recall. For each subject-session, we ranked performance on each list from worst to best, and averaged lists of a given rank across all 23 sessions. In Figure 1B, each subject’s lowest dot thus represents the average, across sessions, of that subject’s recall performance on their “worst” lists. Conversely, each subject’s highest dot represents the average of their recall performance on their “best” lists. Every dot represents an average across 23 lists.

Figure 1 reveals substantial variability in free-recall performance across both sessions and lists. We next sought to identify the sources of this variability. We thus developed a regression model to predict recall based on several likely predictor variables. For our model of intersession variability, we considered four factors: session number (a surrogate for degree of practice at the task), self-reported hours of sleep on the prior night, subjectively-rated alertness, and time-of-day. For our model of inter-list variability, we considered four factors: session number (1-23), block number within a given session (1-3), list number within a block (1-8), and the average recallability of the words on each target list as determined from data obtained from all of the other subjects in the experiment^2^. Table 1 shows the correlation matrix of the predictors for each model. Within each model, the predictors appear to have weak correlations and are thus well-suited for regression analyses.

**Table 1.** *Correlation Matrix of Predictors of Interest for Intersession and Interlist Models*

|  | Session | Sleep | Time | Alertness |
| --- | --- | --- | --- | --- |
| Session | 1.00 | * | * | * |
| Sleep | -0.02 | 1.00 | * | * |
| Time | 0.03 | -0.08 | 1.00 | * |
| Alertness | -0.13 | 0.17 | 0.09 | 1.00 |

**Interlist Predictors**
|  | Session | Block | list | Recallability |
| --- | --- | --- | --- | --- |
| Session | 1.00 | * | * | * |
| Block | 0.00 | 1.00 | * | * |
| list | 0.00 | 0.00 | 1.00 | * |
| Recallability | 0.00 | -0.01 | 0.00 | 1.00 |

The predictors appear to be weakly correlated.

We fit the intersession- and interlist-variability models to data from each of the 79 subjects. Figure 2A illustrates results for the four variables in the intersession-variability model; Figure 2B shows results for the four variables in the interlist-variability model. The dots in each panel indicate the set of subject-specific *β* values for each term in the regression model. Filled circles indicate those *β* values that exceed our significance threshold (*p* < 0.05 FDR corrected for the 79 model fits). As may be seen from the distributions of significant coefficients, some variables exhibited consistent positive or negative effects across subjects (e.g., *block*, *list*, and *recallability* in Panel B), whereas other variables exhibited mixed effects, with some subjects having significant positive coefficients and others having significant negative coefficients (e.g., the *session* variable in Panel A). Although our predictor variables are only weakly correlated, interpretation of regression coefficients may nonetheless be biased. We report these results primarily for illustrative purposes.

**Figure 2.**
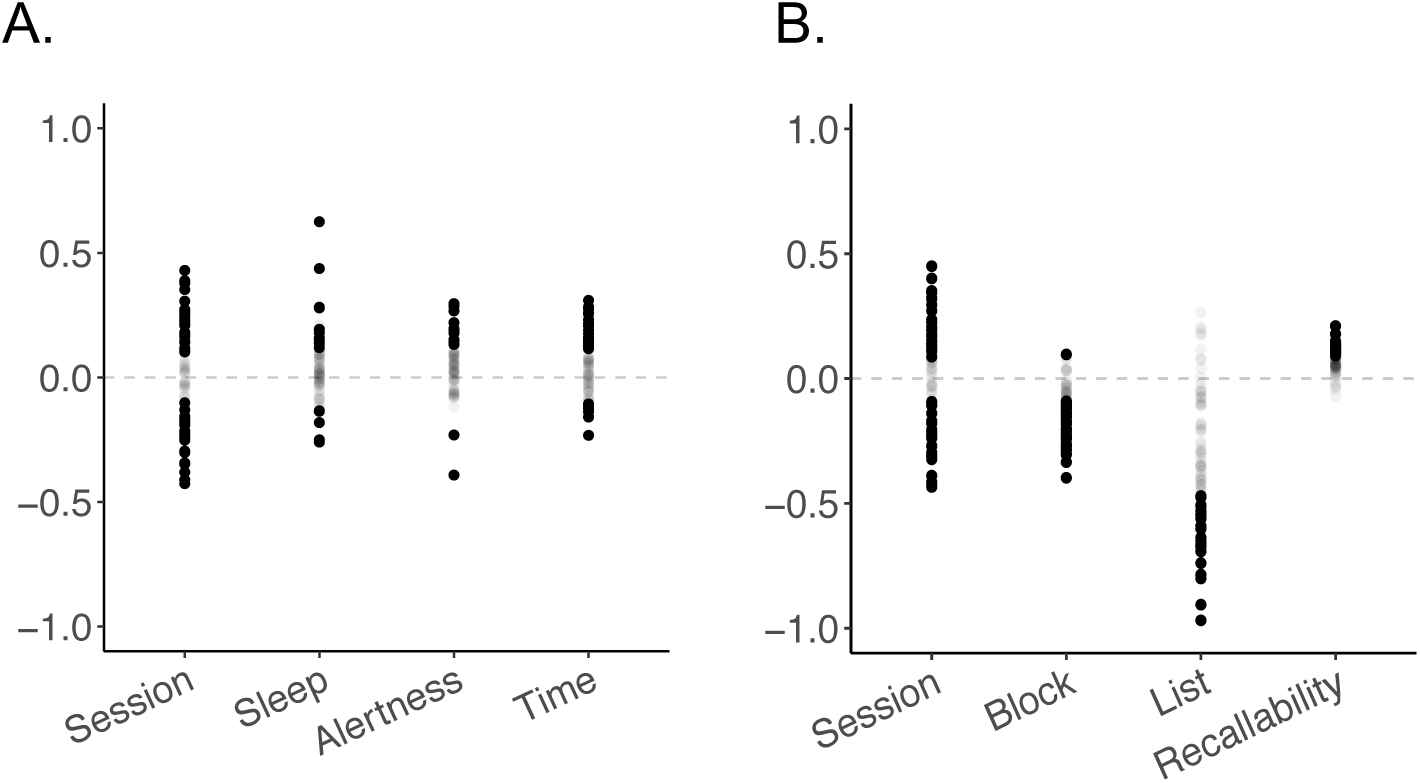
Distributions of beta values for each predictor variable. Each circle denotes the normalized regression coefficient for a single subject, with filled circles indicating coefficients that met an FDR correct *p* < 0.05 significance criterion. Panel A shows the four variables included in the intersession-variability model; Panel B shows the four variables included in the interlist-variability model.

To evaluate which variables were reliably positive or negative across subjects, we utilize linear mixed models (Bates, Mächler, Bolker, & Walker, 2014) that allow for the effects of the predictors to vary across subjects (see Table 2). The subject-level random effects of a predictor are treated as deviations from the fixed effect. As a pre-selection step, we assess whether the effect of each predictor varies across subjects by fitting a linear mixed model with both fixed and random effects on the intercept and the slope of that predictor. Then, the performance of the mixed model is compared to that of a model without random effects via a likelihood ratio test. For the intersession model, all the predictors of interest (session number, amount of sleep, alertness, and start time) vary significantly across subjects and will have both fixed and random effects. For the inter-list model, all predictors also vary significantly across subjects and will have both fixed and random effects. For both models, we logit-transform the response variable (probability of recall for a given list) to remove the range restrictions of probability outcomes.

**Table 2.**
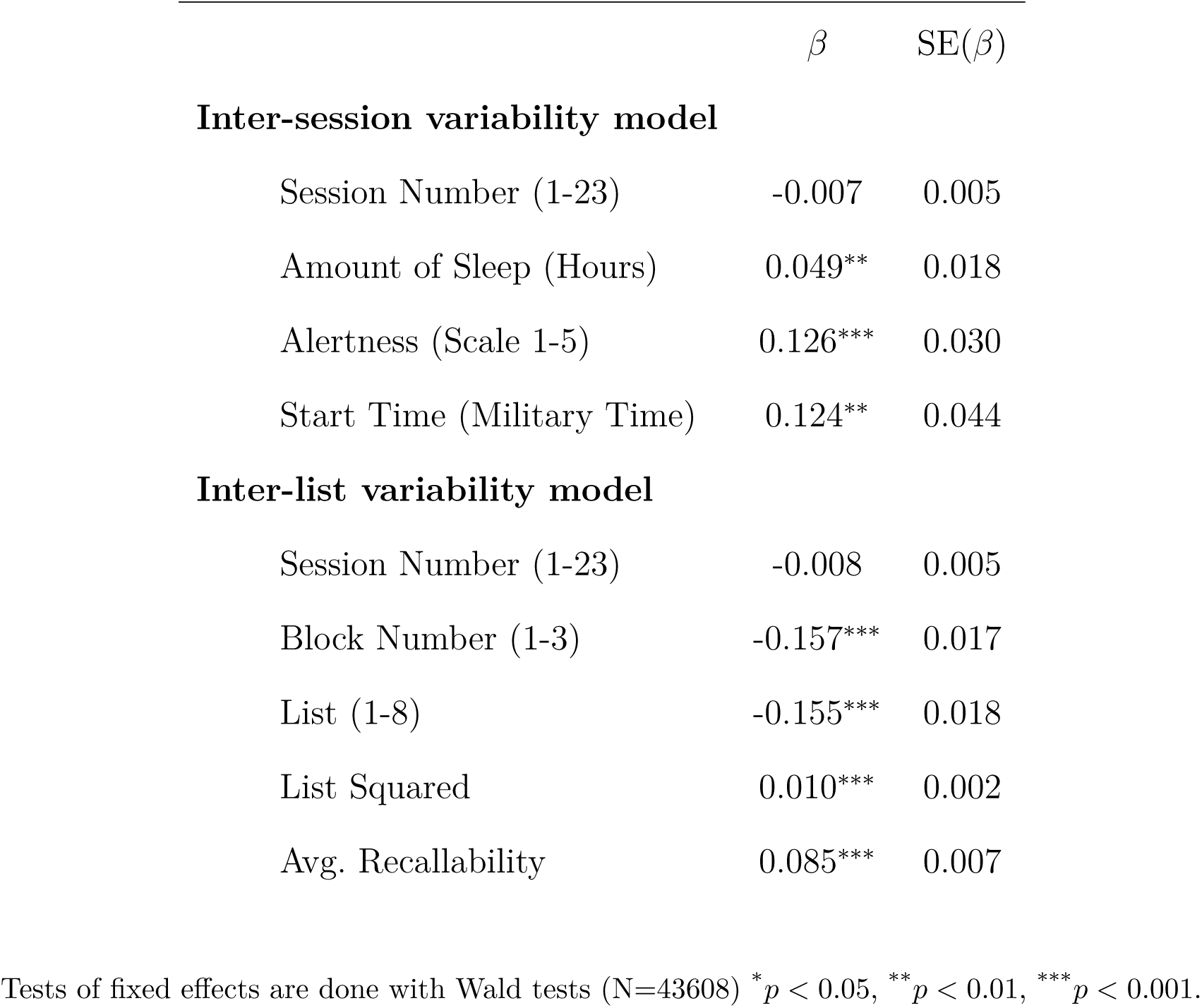
*Fixed Effects of Variables Predicting Probability of Recall*

For our model of intersession variability, *amount of sleep*, *start time*, and *alertness* all exhibit significantly positive effects, indicating that subjects’ recall performance improved when they had a good amount of sleep the night before, when they took the experiment early, and when they were alert before the task. For our model of inter-list variability, both *block number* and *list number* exhibited significant negative effects, indicating that subjects’ recall performance declined over the course of the session. This result is depicted in Figure 3A which shows the change in recall performance across the 24 lists in each session. Because of the non-linear relation between list number and recall performance shown in the Figure, we included a quadratic term in the model (List Squared), which was significantly positive. *Average word recallability* also reliably predicted recall performance across subjects, with lists possessing easier to remember words exhibiting higher overall levels of recall. This is illustrated in Figure 3B which shows the relation between the average recallability of the words in a given list and the actual recall performance for those lists, grouped into 24 bins of recallability.

**Figure 3.**
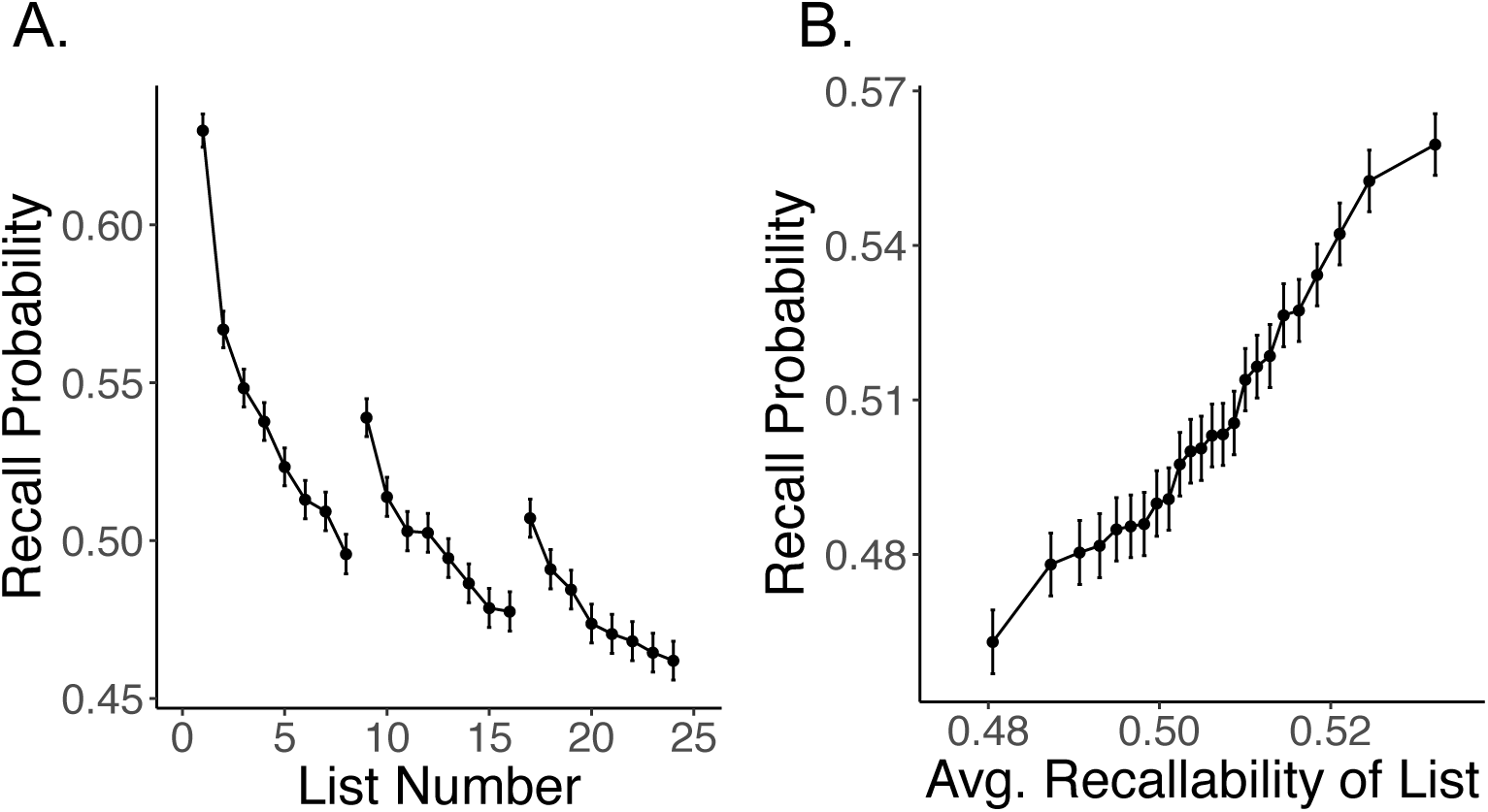
Predictors of interlist variability. **A.** Within each session, recall decreased across successive lists, but increased following the two breaks, consistent with a proactive interference account. **B.** Sorting lists into 24 equally populous bins of predicted recallability we can see that the average recallability of each word within a given list reliably predicts overall list recall. For each subject, recallability was determined based on data from all of the other subjects (see text for details). Error bars indicate ±1*SEM*.

Having modeled inter-session and inter-list variability at the individual subject level, we can now examine the residual variability (i.e., variability not accounted for by our models). Figure 4 shows the residual inter-session and inter-list variability in recall performance, and the degree to which each subject’s model reduced his or her raw variability (shown in Figure 1). Residual values were calculated for the predicted probability of recall for each session (Figure 4A) and for each list (Figure 4B) using the regression models described above, and each of these residuals were added to the subject’s actual mean to produce the resulting adjusted recall distributions. Panels C and D show the percent of the initially-estimated variability reduced by each subject’s best-fitting model.

**Figure 4.**
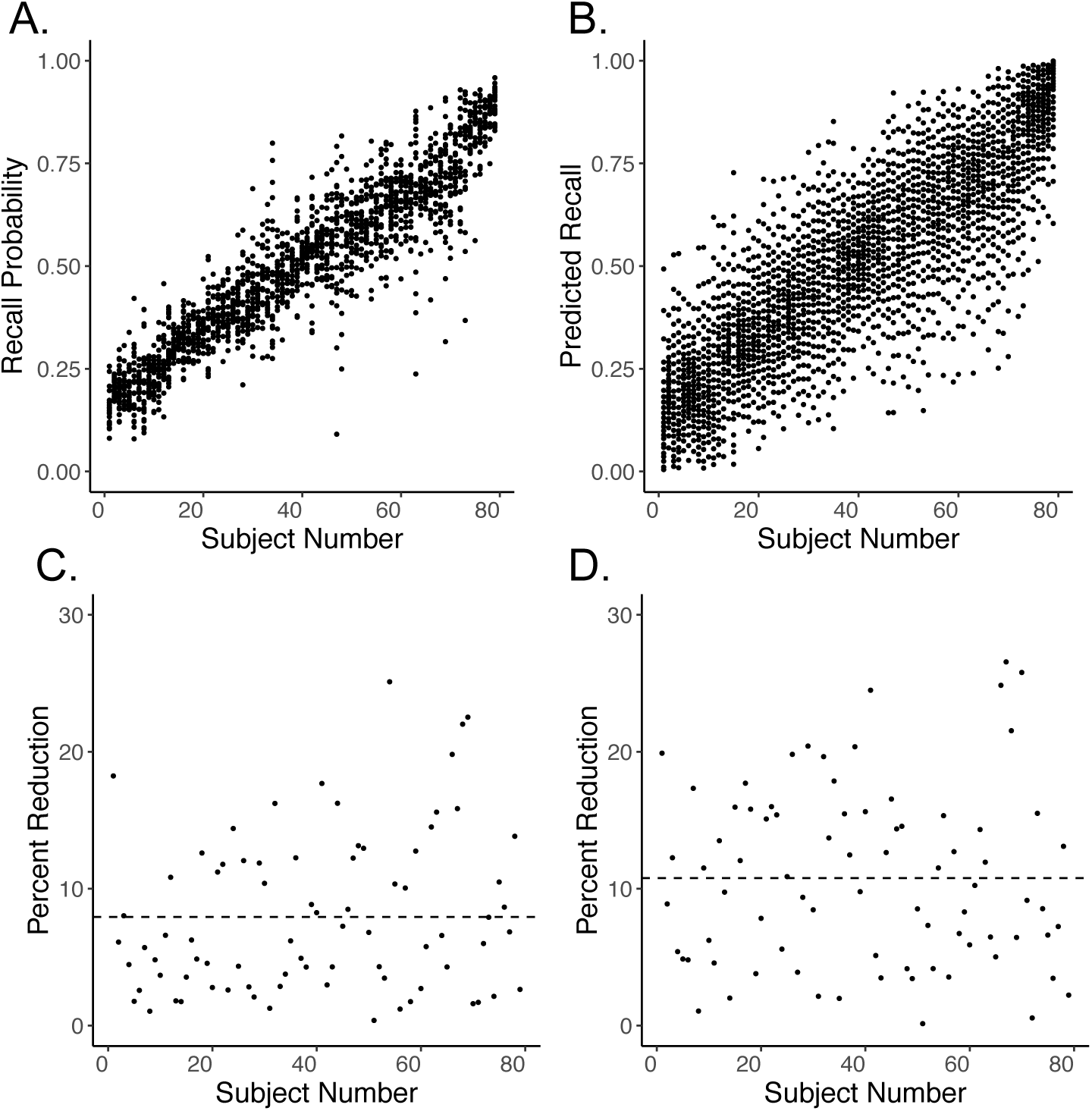
Residual variability in recall. **A. Inter-session variability.** Each dot represents the proportion of words recalled for a single subject in a single session after removing variability accounted for by each subjects predictive model. **B. Inter-list variability.** Each dot here represents the adjusted proportion of words recalled for a single subject on 23 lists, one list taken from each session, ranked by within-session recall performance. In this analysis, recall probability was adjusted for each subject according to his or her predictive model, such that the resulting graphs illustrate the residual variability after accounting for the variables in Table 1. **C., D.** show the percent reduction by the models for each of the subjects. The dashed lines represent the mean percent reductions.

The foregoing analyses estimate the degree of inter-session and inter-list variability in human recall performance. The considerable variability across both sessions and lists likely reflects a wide array of uncontrolled variables, both external and internal to the individual. By fitting a separate regression model to each subject’s 552 study-test lists (or 23 experimental sessions) we were able to quantify the contribution of several important variables to recall performance. In the case of inter-session variability, we considered practice effects, which appeared to account for substantial variability in each subject’s individualized model (with some subjects showing improvements and others showing degradation in performance over sessions). We also considered variability in time of day, the hours of sleep on the prior night and self-rated alertness. In the case of inter-list variability, we considered systematic changes in performance over the course of a session, accounting for both the buildup of proactive interference (PI) across lists, and the release from PI, arising from inter-list breaks. We also considered the recallability of the items in a given list as a surrogate for the effects of item-variables such as word-frequency, concreteness, imageability and the like. A feature of our experiment that made this analysis possible was that each subject received the same set of 576 words across 23 sessions, with each word occurring exactly once per session.

Although substantial variation in performance across both sessions and lists remains unaccounted for by our models, list-level variability appears substantially greater than session-level variability. This should not be too surprising, as session-level data eliminates uncontrolled variability in factors that control list-level difficulty beyond those accounted for by our model. For example, our recallability measure assumes that list difficulty is a function of average item difficulty, when in fact, specific sequences of items can differ in their recallability, and our model does not account for these sequential dependencies.

Although unexplained stimulus variables likely contribute to the observed inter-list variability seen in Figure 4, a comprehensive account of these factors may not suffice in explaining the observed variability. Here we consider the possibility that inter-list variability also reflects endogenous factors, such as stochastic variation in the efficacy of memorial function. Since most stochastic processes exhibit autocorrelations at multiple time scales, one might expect that periods of good memory function are similarly autocorrelated.

To test this endogenous-variability hypothesis, we conducted an autocorrelation analysis on the residuals of the list-level multiple regressions illustrated in Figure 4. After calculating a predicted probability of recall for each subject’s lists, we calculated residuals by subtracting from actual probability of recall for each list. We then applied a linear mixed model with the residual being the response and its lagged-one value being the predictor. We included both random and fixed effects for the predictor. We found that the coefficient for the fixed effect of lagged-one residual is *β* = 0.176 with a standard error of 0.017 (*p* < 0.001), demonstrating a high degree of autocorrelation in the processes giving rise to recall success. We obtained nearly identical results when we included prior-list recall performance as an additional variable in the inter-list linear mixed model described above. In these analyses we were not predicting residual performance, but rather, we simply included all of the variables in a simultaneous regression model. Using this analytic method, we found the fixed effect *β* coefficient of 0.173 with a standard error of 0.017 (*p* < 0.001). Comparing this coefficient with the other predictor variables, we can see that prior-list recall was substantially predictive of current list recall, in support of the endogenous variability account.

## General Discussion

To quantify intra-individual variability in episodic memory we recruited subjects to participate in a 23-session experiment involving study and recall of 552 lists. As shown in Figure 1, subjects exhibited highly variable performance, both within and across sessions. To determine how much of this variability could be explained by known variables, we applied linear mixed effects models to account for both interlist and intersession variability in recall performance. Although each model accounted for significant variability in the recall performance, we were struck by how much variability remained after removing the effects of our predictor variables. The extent to which inter-list variability was reduced, across subjects, ranged from 0.01 to 26.6 percent, with a mean of 11% (± 6.5). Similar results were obtained for the inter-session variability model.

Although our models of inter-list and inter-session variability surely omitted task-related variables that could affect performance, we suspect that much of the residual variability arose not from uncontrolled experimental factors, but rather from endogenous variation in the cognitive processes that support successful memory encoding and retrieval (e.g., Kahana, Rizzuto, & Schneider, 2005). Indeed, recent neuroscientific findings suggest that the brain’s functional connectivity varies stochastically, and that such variation may have important implications for the brain’s computational abilities (Fox, Snyder, Vincent, & Raichle, 2007; He, Zempel, Snyder, & Raichle, 2010; Palva et al., 2013).

Stochastic variability in neural activity supporting memory processes may be expected to exhibit a high degree of temporal autocorrelation (MacDonald, Li, & Bäckman, 2009; MacDonald, Nyberg, & Bäckman, 2006). To test this idea, we examined the behavioral autocorrelation in list-level recall by adding subjects’ recall performance on the prior list as a predictor of current list recall. We found that (residual) prior-list recall was a strong and consistent predictor of (residual) current list recall.

Although our autocorrelation analysis appears consistent with the aforementioned neural results, it is also possible that this reflects variability in the strategies that subjects use to support effective memory encoding and recall. Studies by Battig and colleagues (Battig, 1975, 1979) support the idea that subjects vary their strategies as they learn to perform a task. Although such a strategic variability account is certainly viable, it does not align so well with our finding that the extent of variability appeared fairly consistent across subjects with widely varying overall performance. Furthermore, we did not find any evidence of a decrease in variability across sessions, as might be expected as subjects converge on a consistent strategy for list learning.

How might existing theories of episodic memory account for substantial intra-individual variability in recall performance reported here? Although we have not undertaken a formal analysis of these theories, examination of several successful recall models indicates that the parameters that control encoding and retrieval efficiency are typically assumed to be constant across items, lists, and sessions (Brown, Neath, & Chater, 2007; Farrell, 2012; Kimball, Smith, & Kahana, 2007; Sederberg, Howard, & Kahana, 2008; Sirotin, Kimball, & Kahana, 2005). In models based on retrieved context theory (e.g., Lohnas et al., 2015; Polyn, Norman, & Kahana, 2009), variability in recall would arise due to the idiosyncratic semantic similarity structure of the studied lists, and the probabilistic nature of the retrieval process. Although these mechanisms would give rise to some variability in the number of recalled items across lists (see, e.g., Figure 10 of Polyn et al., 2009) the variability reported here is present even after averaging across many lists.

In fields of study where experimental control is often beyond reach, such as economics, researchers have found that the variance of measured variables provides critical constraints upon theory. For example, stock market volatility appears to be far greater than can be attributed to rational factors, such as the expectation of future dividends (Shiller, 1981). This finding, along with similar results observed for other macro-economic measures, have provided critical constraints on existing theories. Attempts to account for these findings, in turn, have led to important theoretical advances (e.g. Campbell, 2003; Tsai & Wachter, 2015).

It is time for students of memory to move beyond mean performance and consider the role of variability as a source of new knowledge concerning memory. Elevating variability to a variable that we must explain, and even applying functional methods to studying its controllers, will fuel important new insights into our understanding of memory.

## Footnotes

1 Subjects participated in a 24th experimental session during which they studied lists composed of both old words (drawn from the pool of 576) and new words matched on the word attributes. Because the focus of this paper is on variability in performance under constant conditions, our analyses do not include data from this last session.

2 To estimate list recallability, we had considered using a variety of other predictor variables including word frequency, concreteness, imageability, etc. Because each subject in our study saw the same pool of 576 words (24 words x 24 lists per session) in each of 23 sessions, we decided to adopt the more empirical approach of estimating recallability directly based on the aggregate recall data of the other subjects in the experiment. This involved separately computing each word's average recall probability for each subject, based on their 23 encounters with a given word (once in each session). Then, for each subject, we formed an average recallability for each word based on the data obtained across the *N —* 1 remaining subjects in the study.

